# Causal epigenome-wide association study identifies CpG sites that influence cardiovascular disease risk

**DOI:** 10.1101/132019

**Authors:** Tom G. Richardson, Jie Zheng, George Davey Smith, Nicholas J. Timpson, Tom R. Gaunt, Caroline L. Relton, Gibran Hemani

## Abstract

The extent to which genetic influences on complex traits and disease are mediated by changes in DNA methylation levels has not been systematically explored. We developed an analytical framework that integrates genetic fine mapping and Mendelian randomization with epigenome-wide association studies to evaluate the causal relationships between methylation levels and 14 cardiovascular disease traits.

We identified 10 genetic loci known to influence proximal DNA methylation which were also associated with cardiovascular traits (P < 3.83×10^-08^). Bivariate fine mapping suggested that the individual variants responsible for the observed effects on cardiovascular traits at the *ABO*, *ADCY3*, *ADIPOQ, APOA1* and *IL6R* loci were likely mediated through changes in DNA methylation. Causal effect estimates on cardiovascular traits ranged between 0.109-0.992 per standard deviation change in DNA methylation and were replicated using results from large-scale consortia.

Functional informatics suggests that the causal variants and CpG sites identified in this study were enriched for histone mark peaks in adipose tissue and gene promoter regions. Integrating our results with expression quantitative trait loci data we provide evidence that variation at these regulatory regions is likely to also influence gene expression at these loci.

## Introduction

Approximately 88% of trait-associated variants detected using Genome Wide Association Studies (GWAS) reside in non-coding regions of the genome, which suggests that they may be influencing mechanisms which act through gene regulation^1^. Recent studies have incorporated data on genetic variants associated with gene expression (expression quantitative trait loci (eQTL)) into results from GWAS of complex traits to help identify the putative causal variant in a genomic region, as well as provide evidence suggesting which genes may be influenced by this variant^2-5^. This direction of inquiry can be extended to other ‘omic’ data types to gain further insights into the mechanistic pathway between genetic variant and causally associated trait. In this study, we introduce an analytical framework to integrate genetic predictors of DNA methylation levels with complex traits.

DNA methylation is an epigenetic regulation mechanism which has been shown to play a key role in many biological processes and disease susceptibility^6-8^. Recent studies have had success in identifying genetic variants associated with DNA methylation (methylation quantitative trait loci (mQTL)) and report that they appear to overlap with eQTL at a large number of loci across the genome^9; 10^. This suggests that both DNA methylation and gene expression could reside along the causal pathway between genetic variation and disease, although thus far uncovering evidence of a mediated effect between mQTL and traits has been limited in contrast to using eQTL^11-14^. However, identifying epigenetic markers for disease risk should prove valuable in understanding the underlying biological mechanisms for trait-associated variants ^15^.

Mendelian randomization (MR) is a method by which genetic variants robustly associated with modifiable exposures can be used as instrumental variables to infer causality amongst correlated traits ^16; 17^. If DNA methylation resides along the causal pathway between genetic variant and trait, we would expect it to be correlated with our trait of interest. However, much like other traits analysed in epidemiological studies, DNA methylation is prone to confounding and reverse causation. Using an MR framework we can investigate whether DNA methylation has a causal relationship with a phenotypic outcome, suggesting that it may reside along the causal pathway to disease ^18^. Moreover, as discussed in a recent review, MR has advantages over alternative approaches in mediation analysis (such as the Causal Inference Test ^19^), as it can detect the correct direction of effect in the presence of measurement error ^20^.

Recent approaches to MR have shown that the robustness of causal inference is improved if there are many instruments because one can evaluate whether the SNP effects on the causal trait are proportional to the SNP effects on the consequential trait ^16; 21^. We exploit this property to evaluate the causal influence of complex traits (which typically have many instruments) on DNA methylation (i.e. bi-directional MR^22^). But a pitfall of evaluating the causal influence of DNA methylation on complex traits is that DNA methylation is typically instrumented by only a single cis-acting variant. Hence, an unreliable MR estimate of causality could arise due to the mQTL simply being in linkage disequilibrium with a variant that influences the cardiovascular trait through means other than the methylation level.

There are methods which have been devised to address this issue for eQTL within a two-sample framework or using multiple instruments ^2; 23-25^, although thus far there are no appropriate methods when effect estimates are obtained using the same sample and where only one valid instrument exists. Therefore, to distinguish linkage disequilibrium from mediation we integrate fine mapping to evaluate the likelihood of the mQTL being the same causal variant as the SNP influencing the cardiovascular trait. We have also undertaken functional informatics and incorporated eQTL data as this may support findings suggesting that DNA methylation resides on the causal pathway between variant and disease. However, a limitation of using single variant instruments in general is that it is not possible to reliably distinguish horizontal pleiotropy from mediation ^26^.

Together, the causal relationships between DNA methylation and cardiovascular traits are delineated into four potential categories (Figure 1):

**Figure 1:**
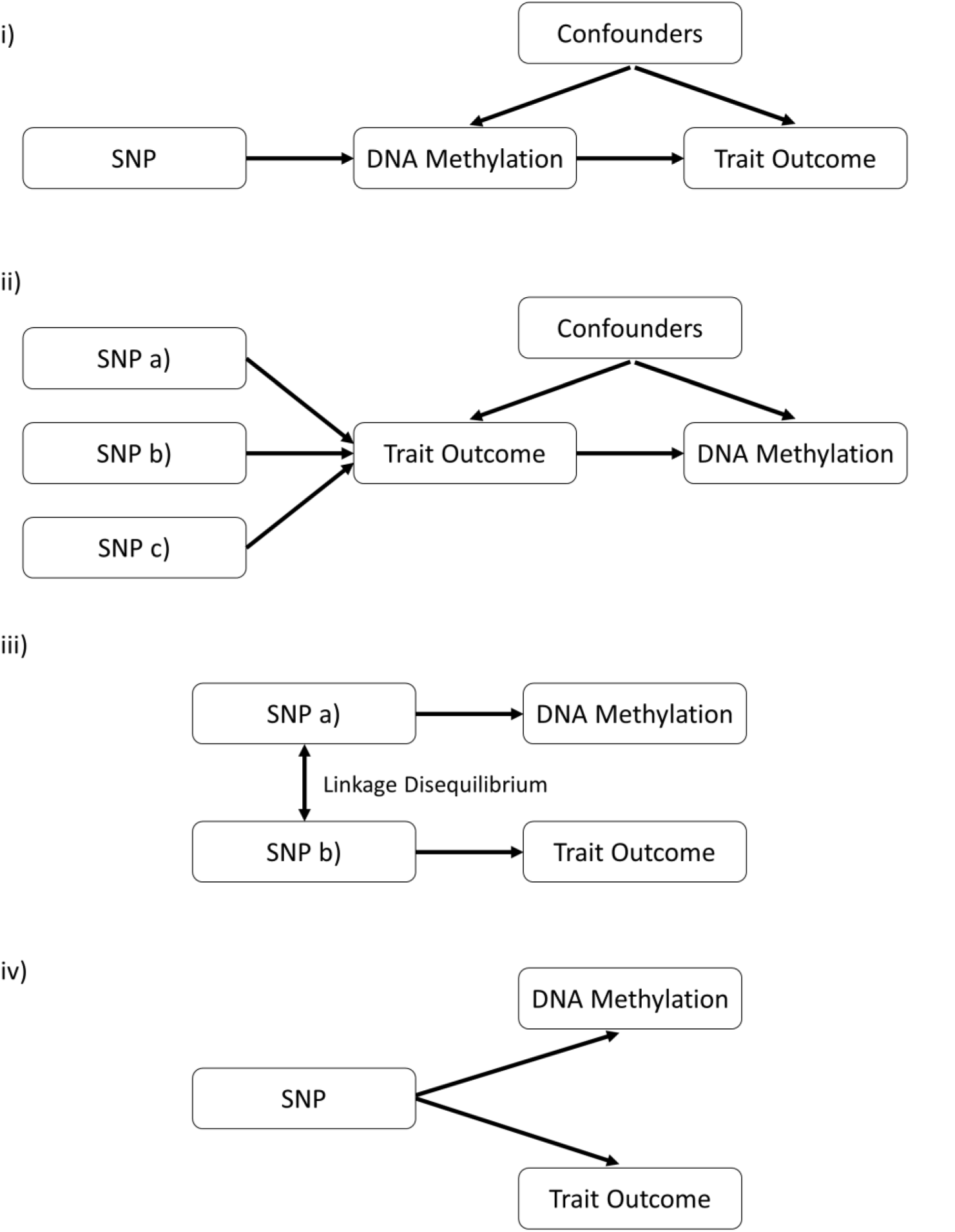
Explanations evaluated to explain observed associations between methylation quantitative trait loci and trait outcomes. i) The genetic variant has an effect on the phenotype, mediated through DNA methylation. ii) The genetic variant has an effect on the phenotype by alternative biological mechanisms, which then has a downstream effect on DNA methylation at this locus. iii) The genetic variant which influences DNA methylation is simply in linkage disequilibrium with another variant which is influencing the associated trait. iv) The genetic variant is influencing both DNA methylation and phenotype by two independent biological pathways (also known as horizontal pleiotropy).

1. The genetic variant has an effect on the phenotype, mediated by DNA methylation.
2. The genetic variant has an effect on the phenotype by alternative biological mechanisms, which then has a downstream effect on DNA methylation at this locus.
3. The genetic variant which influences DNA methylation is simply in linkage disequilibrium (LD) with another variant which is influencing the associated trait.
4. The genetic variant is influencing both DNA methylation and phenotype by two independent biological pathways (also known as horizontal pleiotropy).

We have developed a framework to systematically navigate through these scenarios and have applied it to analyse 14 different cardiovascular traits. In our discovery analysis we used genotype and DNA methylation data from prepubertal individuals to discover causal pathways on early childhood phenotypes. Replication was then undertaken using GWAS summary statistics from large-scale consortia.

## Materials and Methods

### The Avon Longitudinal Study of Parents and Children (ALSPAC)

ALSPAC is a population-based cohort study investigating genetic and environmental factors that affect the health and development of children. The study methods are described in detail elsewhere ^27; 28^ (http://www.bristol.ac.uk/alspac). Briefly, 14,541 pregnant women residents in the former region of Avon, UK, with an expected delivery date between 1^st^ April 1991 and 31^st^ December 1992, were eligible to take part in ALSPAC. Detailed information and biosamples have been collected on these women and their offspring at regular intervals, which are available through a searchable data dictionary (http://www.bris.ac.uk/alspac/researchers/data-access/data-dictionary/).

Written informed consent was obtained for all study participants. Ethical approval for the study was obtained from the ALSPAC Ethics and Law Committee and the Local Research Ethics Committees.

### Accessible Resource for Integrative Epigenomic Studies project (ARIES)

#### Samples

Blood samples were obtained for 1,018 ALSPAC mother-offspring pairs (mothers at two timepoints and their offspring at three timepoints) as part of the Accessible Resource for Integrative Epigenomic Studies project (ARIES)^29^. The Illumina HumanMethylation450 (450K) BeadChip array was used to measure DNA methylation at over 480,000 sites across the epigenome.

#### Methylation assays

DNA samples were bisulfite treated using the Zymo EZ DNA Methylation^TM^ kit (Zymo, Irvine, CA). The Illumina HumanMethylation450 BeadChip (HM450k) was used to measure methylation across the genome and the following arrays were scanned using Illumina iScan, along with an initial quality review using GenomeStudio. A purpose-built laboratory information management system (LIMS) was responsible for generating batch variables during data generation. LIMS also reported quality control (QC) metrics for the standard probes on the HM450k for all samples and excluded those which failed QC. Data points with a read count of 0 or with low signal:noise ratio (based on a p-value > 0.01) were also excluded based on the QC report from Illumina to maintain the integrity of probe measurements. Methylation measurements were then compared across timepoints for the same individual and with SNP-chip data (HM450k probes clustered using k-means) to identify and remove sample mismatches. All remaining data from probes was normalised with the Touleimat and Tost^30^ algorithms using R with the wateRmelon package^31^. This was followed by rank-normalising the data to remove outliers. Potential batch effect were removed by regressing data points on all covariates. These included the bisulfite-converted DNA (BCD) plate batch and white blood cell count which was adjusted for using the *estimateCellCounts* function in the minfi Bioconductor package^32^.

#### Genotyping assays

Genotype data were available for all ALSPAC individuals enrolled in the ARIES project, which had previously undergone quality control, cleaning and imputation at the cohort level. ALSPAC offspring selected for this project had previously been genotyped using the Illumina HumanHap550 quad genome-wide SNP genotyping platform (Illumina Inc, San Diego, USA) by the Wellcome Trust Sanger Institute (WTSI, Cambridge, UK) and the Laboratory Corporation of America (LCA, Burlington, NC, USA). Samples were excluded based on incorrect sex assignment; abnormal heterozygosity (<0.320 or >0.345 for WTSI data; <0.310 or >0.330 for LCA data); high missingness (>3%); cryptic relatedness (>10% identity by descent) and non-European ancestry (detected by multidimensional scaling analysis). After QC, 500,527 SNP loci were available for the directly genotype dataset. Following QC the final directly genotyped dataset contained 526,688 SNP loci.

#### Imputation

Imputation was performed using a joint reference panel using variants discovered through whole genome sequencing (WGS) in the UK10K project ^33^ along with known variants taken from the 1000 genomes reference panel. Novel functionality was developed in IMPUTE2 ^34^ to use each reference panel to impute missing variants in their counterparts before ultimately combining them together. All variants were filtered to have Hardy-Weinberg equilibrium P > 5×10^-7^ and an imputation quality score ≥ 0.8 or higher.

#### Phenotypes

ALSPAC individuals were measured for 14 different cardiovascular traits. These were body mass index (**BMI**), systolic blood pressure (**SBP),** diastolic blood pressure (**DBP**), total cholesterol (**TC**), triglycerides (**TG**), high density lipoprotein cholesterol (**HDL**), low density lipoprotein cholesterol (**LDL**), apolipoprotein A (**Apo A1),** apolipoprotein b (**Apo B)**, interleukin 6 (**IL-6**), adiponectin, C-reactive protein (**CRP**) and leptin. Methods for phenotyping can be found in the supplementary material.

### Statistical Analysis

We undertook a methylome-wide association study (MWAS) to evaluate the association between all eligible mQTL and each trait in turn. This was decided over a conventional epigenome-wide association study (EWAS) (i.e. evaluating the association between methylation levels at CpG sites and traits) due to a larger proportion of individuals in ALSPAC having genotype data rather than 450K data after merging on phenotypes.

All variants with previous evidence of genetic association with DNA methylation in ARIES (referred to hereafter as mQTL) were eligible for analysis ^8^. We used a strict threshold to define mQTL (P<1.0×10^-14^) as we were unable to identify a suitable replication cohort for observed effects on methylation levels derived from blood. Furthermore, this strict threshold reduces the risk of MR analyses suffering from weak instrument bias. We also removed mQTL associated with 450K probes flagged for exclusion based on evaluations by Naeem et al ^35^, based on their criteria of overlapping SNPs at CpG probes, probes which map to multiple locations and repeats on the 450K array. Moreover, we excluded mQTL associated with a CpG site which was over 1Mb distance away (known as trans-mQTL), therefore leaving mQTL which were only associated with a nearby CpG site (known as cis-mQTL). This was to reduce the possibility of pleiotropy in our analysis as variants which associated with methylation at multiple CpG sites across the epigenome may be influencing independent biological pathways simultaneously. However, we cannot rule out the possibility that in future studies that the mQTL included in this study are in fact influencing multiple CpG sites, although we do not currently have an adequate sample size to detect these additional trans effects.

This was important to include in our study design as we anticipated that single instrument MR analyses may be necessary at a later stage when evaluating causal effects, which therefore restricts our ability to investigate pleiotropy using multiple valid instruments. GCTA was used to undertake a conditional analysis and determine independent loci for each CpG site ^36^. mQTL were analysed in turn with each trait using linear regression with adjustment for age and sex. Results were plotted using a Manhattan plot using code derived from the qqman R package^37^. Scripts to generate this plot are available at https://github.com/MRCIEU/qqman_multiple_colours.

### Mendelian randomization analysis

Observed associations between genotype and traits which survived a stringent multiple testing threshold (i.e. P < 0.05/number of tests undertaken) were then analysed using Mendelian randomization (MR) to discern whether a causal effect existed of DNA methylation on cardiovascular traits. This was undertaken using two stage least squares (2SLS) regression with DNA methylation as our exposure, phenotypic trait as our outcome and using the relevant mQTL as our instrumental variable. Measures of DNA methylation were initially taken from the childhood time point in ARIES (mean age: 7.5, standard deviation: 0.15) as this was the closest time point to phenotype measurements. Follow-up analyses were also undertaken using methylation data from the birth time point (using cord blood) and the adolescent time point (mean age: 17.1, standard deviation: 1.01). The R package ‘systemfit’^38^ was used to obtain causal effect estimates using two-stage least squares.

We replicated observed effects by undertaking a two sample MR analysis (2SMR)^39^ using estimated effects between genetic variants and associated traits obtained from published studies. Moreover, a two-sample framework removes any potential bias encountered in the discovery analysis due to effects on both methylation and trait being obtained in the same sample. When observed effects for sentinel mQTL were not available from published studies we used variants in LD with these SNPs instead (r^2^ > 0.8).

Figure 1 illustrates the 4 possible explanations investigated where evidence of a causal effect was observed using MR. Figure 2 provides an overview of our approach to investigate these explanations. To robustly test explanation ii) we performed the reverse MR analysis, evaluating if the cardiovascular trait influenced the DNA methylation level. Instruments for this analysis were identified using the NHGRI-EBI GWAS catalog^40^. Relevant GWAS for interleukin-6 were not available at the time of analysis and so we identified instruments based on findings from Naitza et al ^41^ (P < 5.0×10^- 08^). A lack of evidence suggesting a causal relationship in this analysis would suggest that explanation ii) was unlikely in each instance.

**Figure 2:**
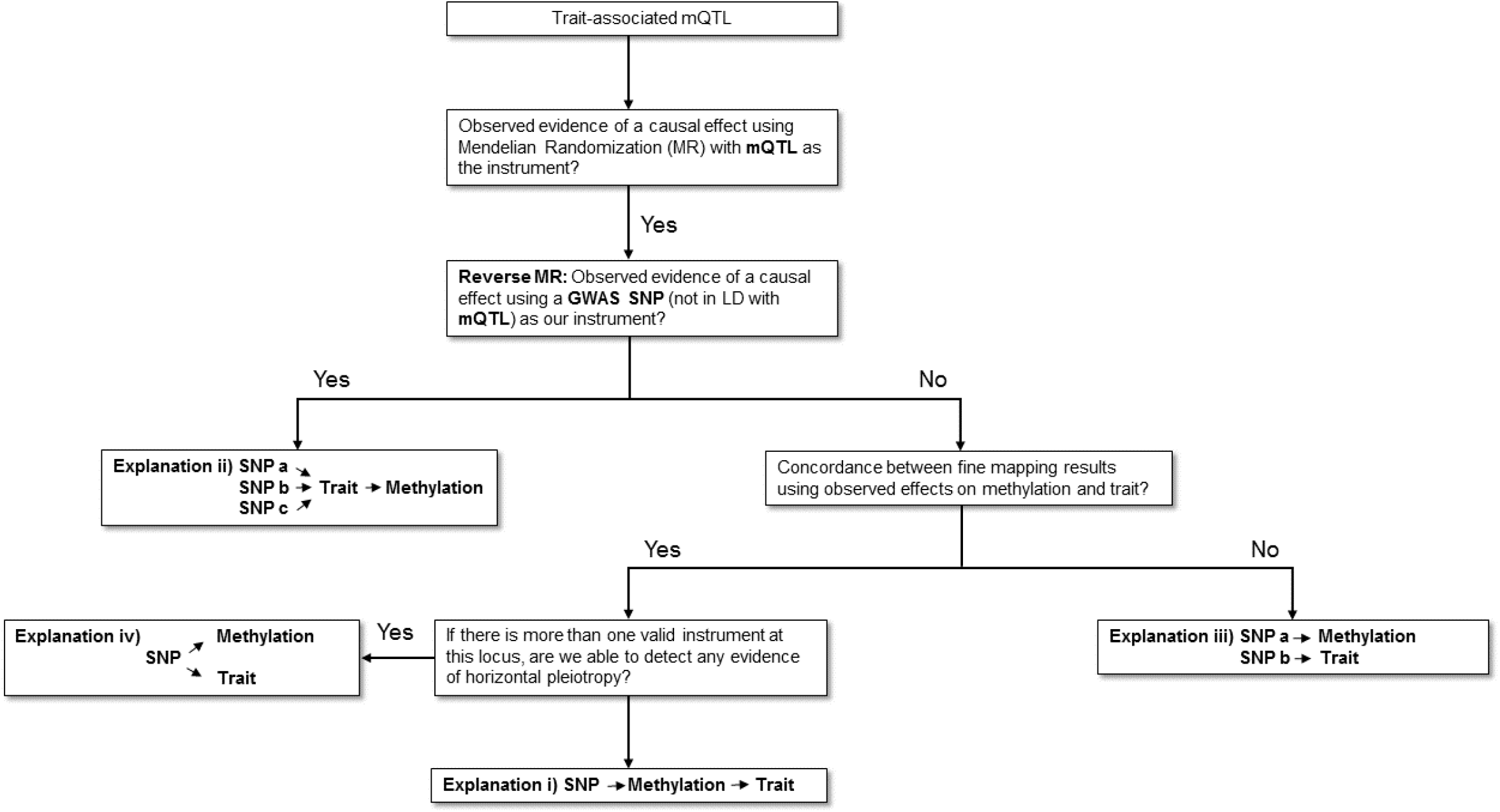
Analysis pipeline to evaluate explanations for observed associations between methylation quantitative trait loci and trait outcomes. This flowchart provides an overview of the analysis plan in this study to evaluate 4 different explanations which may explain trait-associated methylation quantitative trait loci (mQTL). LD – linkage disequilibrium, GWAS – Genome-wide association study.

**Figure 3:**
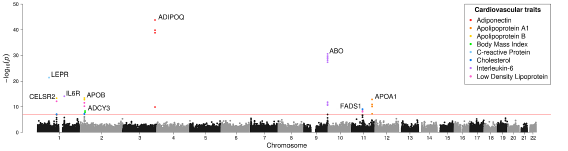
Manhattan plot illustrating observed association between methylation quantitative trait loci and cardiovascular traits. Manhattan plot illustrating the observed association between methylation quantitative trait loci (mQTL) and various cardiovascular traits. Points represent – log10 p-values (y-axis) for genetic variants according to their genomic location (x-axis). Effects that survive the multiple testing threshold in our analysis (P < 9.45×10^-08^ – represented by the red horizontal line) are coloured according to their associated trait and annotated according to the likely impacted gene.

### Bivariate fine mapping

Bivariate fine mapping was undertaken using FINEMAP^42^ at each locus detected in the previous analysis. FINEMAP generates a Bayes factor for each variant at a locus which reflects the likelihood that it is the underlying causal variant at this region. Bivariate fine mapping requires all variants at a locus to be fine mapped using two different effect estimates 1) observed effects between SNPs and DNA methylation and 2) observed effects between SNPs and outcome phenotypes. Estimates between all variants in high LD (r^2^ ≥ 0.8) with the sentinel SNP for each association signal were used for bivariate fine mapping analyses.

This analysis was undertaken to evaluate explanation iii), that the mQTL analysed may simply be in LD with the putative causal variant for the phenotypic trait. This was necessary as when evaluating the relationship between DNA methylation at a CpG site and outcome trait there may likely only be one valid instrumental variable (i.e. the mQTL at this region). Bivariate fine mapping in this instance therefore evaluates whether the causal mQTL at a locus is likely the same causal variant for the observed effect on outcome trait. However, it does not rule out the possibility of a single variant influencing DNA methylation and outcome trait through independent biological pathways (i.e. explanation iv).

Concordance between the top SNPs for the two sets of fine mapping analyses would suggest that explanation i) may be responsible for the observed effect and that DNA methylation resides on the causal pathway between variant and phenotypic trait. Bivariate fine mapping using effect estimates for both methylation and cardiovascular traits was advantageous in this study as we were able to obtain estimates for all SNPs in our data set without having to rely on summary statistics. To further evaluate explanation iii), we also used the joint likelihood mapping (JLIM) approach^23^. Although JLIM doesn’t specify the likely causal variant at a region, it can be used to examine whether the underlying causal variation is responsible for observed effects on both methylation and cardiovascular trait in a two-sample framework.

### Impact of mQTL on gene expression and histone modification

We applied 2SMR to evaluate the relationship between methylation and expression using observed effects between SNPs and expression in relevant tissue types from the GTEx consortium^43^. When observed effects for sentinel mQTL were not available from GTEx we identified a surrogate SNP instead (r^2^ > 0.8).

We also assessed whether any mQTL were in LD (r^2^ > 0.8) with any previously reported histone quantitative trait loci (hQTL)^44^. When this was true, we applied 2SMR to evaluate the causal relationship between methylation and histone modification at these loci. This analysis was for exploratory purposes as there are aspects of the relationship between DNA methylation and histone modification which remain unexplored, despite progress by recent studies^45; 46^.

### Functional informatics

The variant effect predictor (VEP)^47^ was applied to the top ranked mQTL from the bivariate fine mapping analysis to calculate their predicted consequence. Regulatory data was obtained from Ensembl^48^ to evaluate whether mQTL and CpG sites were located within regulatory regions of the genome. Additional regulatory information was also obtained from the 450K annotation file from Illumina. As we were interested in cardiovascular and lipid traits in this study, adipose tissue data from the Roadmap Epigenomics Project^49^ was used to infer whether the potential causal variants and CpG sites at each locus resided within histone mark peaks.

Enrichment analysis was undertaken to test whether lead SNPs and associated CpG sites were located in regulatory regions more than can be accounted for by chance. To calibrate background expectations, we obtained matched SNPs using snpSNAP^50^ and identified matched CpG sites by randomly sampling probes from the 450K array which were in similar regions across the genome (i.e. within CpG islands/1^st^ Exons etc.). Enrichment was investigated using the hypergeometric test and multiple testing was accounted for by randomly sampling controls SNPs/probes and re-running analyses for 10,000 iterations.

## Results

### Mining for causal influences of methylation on cardiovascular traits

We undertook 529,368 tests to evaluate the association between previously identified mQTL in ARIES with each trait in turn (37,812 unique variants×14 traits). We identified 10 association signals which, after multiple testing correction, provided strong evidence of association (P < 9.45×10^-08^ (i.e. 0.05/529,368)) and can be found in Table 1. The 10 sentinel mQTL identified in this analysis were only strongly associated with DNA methylation at a proximal CpG site and not any other CpG sites in the epigenome based on our findings in ARIES.

**Table 1:** Results of linear regression analysis between methylation trait loci (mQTL) and traits – MAF In these tables

| SNP | Gene | CpG | Trait | Sample Size | Beta | SE | P-value | % explained |
| --- | --- | --- | --- | --- | --- | --- | --- | --- |
| rs266772 | *ADIPOQ* | cg05578595 | Adiponectin | 4248 | -0.992 | 0.070 | $1.72 \times 10^{-44}$ | 4.51% |
| rs687621 | *ABO* | cg21160290 | Interleukin-6 | 4241 | -0.265 | 0.022 | $1.15 \times 10^{-31}$ | 3.05% |
| rs13375019 | *LEPR* | cg04111102 | C-reactive protein | 4251 | -0.213 | 0.022 | $2.65 \times 10^{-22}$ | 2.20% |
| rs7549250 | *IL6R* | cg02856953 | Interleukin-6 | 4241 | -0.176 | 0.022 | $9.71 \times 10^{-16}$ | 1.40% |
| rs169109 | *ADIPOQ* | cg05578595 | Adiponectin | 4248 | -0.167 | 0.022 | $1.44 \times 10^{-14}$ | 1.34% |
| rs541041 | *APOB* | cg25035485 | Apo B | 4251 | -0.209 | 0.028 | $3.76 \times 10^{-14}$ | 1.32% |
| rs7528419 | *CELSR2* | cg00908766 | Apo B | 4251 | -0.196 | 0.026 | $4.63 \times 10^{-14}$ | 1.30% |
| rs625145 | *APOA1* | cg04087571 | Apo A1 | 4251 | 0.200 | 0.027 | $9.78 \times 10^{-14}$ | 0.94% |
| rs174544 | *FADS1* | cg19610905 | Cholesterol | 4250 | -0.143 | 0.023 | $8.61 \times 10^{-10}$ | 0.86% |
| rs6749422 | *ADCY3* | cg01884057 | Body mass index | 6076 | 0.109 | 0.018 | $1.28 \times 10^{-9}$ | 0.55% |

- SNP – Single Nucleotide Polymorphism, Gene – likely implicated gene, CpG – 450K probe ID, Trait – Associated Trait, Sample Size – sample size for this effect, Beta – Observed effect size (units in standard deviations), SE – Standard Error of the effect size, P-value – P-value for observed effect, % explained – proportion of variance in trait explained by mQTL.

### Inferring causal relationships

Causal effect estimates between methylation and cardiovascular traits were obtained at each locus in the MR analysis using mQTL as our instrumental variables (Table 2). Taking these putative associations forward, we evaluated the potential for reverse causal relationships by performing MR of the cardiovascular traits against the DNA methylation levels using SNPs from GWAS as our instruments. There was no evidence to suggest that the putative associations were due to the cardiovascular traits influencing the methylation levels (Supplementary Tables 1) and therefore suggests that these effects can be attributed to explanation i) rather than explanation ii) at these loci (i.e. variation in DNA methylation is responsible for changes in cardiovascular function).

**Table 2:** Results of Mendelian randomization analysis between DNA methylation and traits

| **SNP** | **Gene** | **CpG** | **Trait** | **Sample Size** | **Beta** | **SE** | **P-value** |
| --- | --- | --- | --- | --- | --- | --- | --- |
| rs266772 | *ADIPOQ* | cg05578595 | Adiponectin | 646 | -0.846 | 0.168 | 5.93 x 10<sup>-7</sup> |
| rs687621 | *ABO* | cg21160290 | Interleukin-6 | 646 | -0.293 | 0.061 | 1.77 x 10<sup>-6</sup> |
| rs13375019 | *LEPR* | cg04111102 | C-reactive protein | 646 | -0.265 | 0.076 | 0.001 |
| rs7549250 | *IL6R* | cg02856953 | Interleukin-6 | 646 | 0.468 | 0.175 | 0.008 |
| rs169109 | *ADIPOQ* | cg05578595 | Adiponectin | 646 | -0.363 | 0.121 | 0.003 |
| rs541041 | *APOB* | cg25035485 | Apo B | 646 | 0.298 | 0.114 | 0.009 |
| rs7528419 | *CELSR2* | cg00908766 | Apo B | 646 | 0.271 | 0.064 | 2.74 x 10<sup>-5</sup> |
| rs625145 | *APOA1* | cg04087571 | Apo A1 | 646 | -0.301 | 0.082 | 2.68 x 10<sup>-4</sup> |
| rs174544 | *FADS1* | cg19610905 | Cholesterol | 646 | -0.363 | 0.121 | 0.003 |
| rs6749422 | *ADCY3* | cg01884057 | Body mass index | 846 | 0.106 | 0.048 | 0.028 |

- SNP – Single Nucleotide Polymorphism, Gene – likely implicated gene, CpG – 450K probe ID, Trait – Associated Trait, Sample Size – sample size for this effect, Beta – Observed effect size (units in standard deviations), SE – Standard Error of the effect size, P-value – P-value for observed effect

Using methylation data from 2 other time points across the life course (at birth and adolescence (mean age: 17.1)) we observed consistent directions of effect as was observed using data from the childhood time point (mean age: 7.5) (Supplementary Tables 2 & 3). Evidence of association was observed at each locus in this analysis except for the *ABO* and *IL6R* loci when using the cord data.

We reproduced similar effects for 9 of the 10 mQTLs on cardiovascular traits using effect estimates from published studies (Table 3). The only locus we were not able to find a replication effect estimate for was the mQTL at *IL6R* as it was not in LD (r^2^ > 0.8) with any previously published findings for interleukin 6. Effect estimates suggested a direct relationship between methylation and cardiovascular traits at the *IL6R*, *APOB*, *CELSR2* and *ADCY3* loci (i.e. increased methylation results in an observed increase in the cardiovascular trait), whereas an inverse relationship was observed at the *ADIPOQ*, *ABO*, *LEPR*, *APOA1* and *FADS1* loci (i.e. increased methylation causes a decrease in cardiovascular trait levels).

**Table 3:** Results of replication analysis using two-sample Mendelian randomization

| SNP | Gene | CpG | CpG effect | Trait effect | 2SMR effect | P-value | Study |
| --- | --- | --- | --- | --- | --- | --- | --- |
| rs266772 | *ADIPOQ* | cg05578595 | 0.982 (0.103) | -0.629 (0.143) | -0.641 (0.160) | $6.50 \times 10^{-5}$ | UK10K Consortium (TwinsUK individuals only) (2015) |
| rs687621 | *ABO* | cg21160290 | 0.912 (0.036) | -0.245 (0.026) | -0.269 (0.03) | $9.16 \times 10^{-19}$ | Naitza et al (2012) |
| rs2211651* | *LEPR* | cg04111102 | 0.682 (0.036) | -0.170 (0.022) | -0.249 (0.035) | $3.09 \times 10^{-13}$ | Reiner et al (2012) |
| rs541041 | *APOB* | cg25035485 | 0.627 (0.053) | 0.098 (0.013) | 0.156 (0.025) | $2.05 \times 10^{-10}$ | Kettunen et al (2016) |
| rs169109 | *ADIPOQ* | cg05578595 | 0.383 (0.036) | -0.052 (0.005) | -0.136 (0.017) | $2.58 \times 10^{-15}$ | Dastani et al (2013) |
| rs7528419 | *CELSR2* | cg00908766 | -0.980 (0.037) | -0.089 (0.012) | 0.091 (0.013) | $9.20 \times 10^{-13}$ | Kettunen et al (2016) |
| rs625145 | *APOA1* | cg04087571 | -0.884 (0.044) | 0.057 (0.013) | -0.064 (0.015) | $1.84 \times 10^{-5}$ | Kettunen et al (2016) |
| rs174544 | *FADS1* | cg19610905 | -0.655 (0.031) | 0.047 (0.004) | -0.072 (0.007) | $9.73 \times 10^{-25}$ | Global Lipids Genetics Consortium (2013) |
| rs6749422 | *ADCY3* | cg01884057 | 0.908 (0.026) | 0.068 (0.007) | 0.075 (0.008) | $8.05 \times 10^{-21}$ | Felix et al (2016) |

- \* indicates surrogate variant used ( $r^2 > 0.8$ ), SNP – Single Nucleotide Polymorphism, Gene – likely implicated gene, CpG – 450K probe ID, CpG effect – effect estimate of SNP on methylation, Trait effect – effect estimate of SNP on trait, 2SMR effect – effect estimates from 2-Sample MR analysis, P-value – P-value for observed effect, Study – published study where effect estimates for traits were obtained.

### Evaluating causal variants to infer mediated effects

There was concordance amongst the top SNPs in the bivariate fine mapping analyses at the *ABO*, *ADCY3* and *ADIPOQ* (common signal) loci, as the variant with the largest Bayes factor was the same for the effect on DNA methylation and outcome trait (Supplementary Tables 4). These results lend support to the hypothesis that DNA methylation resides on the causal pathway between genetic variant and outcome trait (i.e. explanation i). At the *APOA1* and *IL6R* loci, the variant with the largest Bayes factor based on effect estimates on DNA methylation had the second largest Bayes factor for observed effect on outcome traits. There was a lack of concordance for the results at the *ADIPOQ* (low frequency signal), *APOB*, *CELSR2*, *FADS1* and *LEPR* loci, suggesting that the mQTL may be in LD with the putative causal variant for the phenotypic trait (i.e. explanation iii). Results of the JLIM method supported evidence at the *ADIPOQ*, *ADCY3* & *APOA1* loci, although we were unable to further evaluate signals at the *ABO* & *IL6R* regions due to unavailable GWAS summary results for interleukin-6 (Supplementary Table 5).

**Table 4:** Results of analysis investigating causal relationship between methylation and expression using Two-Sample Mendelian randomization

| SNP | Gene | CpG | CpG effect | eQTL effect | eQTL P-value | eQTL tissue | 2SMR | P-value |
| --- | --- | --- | --- | --- | --- | --- | --- | --- |
| rs116552240* | *ABO* | cg21160290 | 0.912 (0.036) | 0.548 (0.069) | $1.316 \times 10^{-13}$ | Adipose | 0.601 (0.079) | $3.28 \times 10^{-14}$ |
| rs6737082 | *ADCY3* | cg01884057 | 0.908 (0.026) | 0.208 (0.047) | $1.456 \times 10^{-5}$ | Adipose | 0.229 (0.052) | $1.13 \times 10^{-5}$ |
| rs266772 | *ADIPOQ* | cg05578595 | 0.982 (0.103) | -0.339 (0.078) | $1.893 \times 10^{-5}$ | Adipose | -0.345 (0.087) | $7.67 \times 10^{-5}$ |
| rs688456 | *APOA1* | cg04087571 | -0.884 (0.044) | 0.420 (0.095) | $1.789 \times 10^{-5}$ | Heart | -0.475 (0.11) | $1.58 \times 10^{-5}$ |
| rs541041 | *APOB* | cg25035485 | -0.627 (0.053) | -0.370 (0.066) | $6.326 \times 10^{-8}$ | Heart | 0.590 (0.116) | $4.06 \times 10^{-7}$ |
| rs646776 | *CELSR2* | cg00908766 | -0.980 (0.037) | -1.11 (0.117) | $2.463 \times 10^{-15}$ | Liver | 1.133 (0.127) | $4.20 \times 10^{-19}$ |
| rs174559 | *FADS1* | cg19610905 | -0.655 (0.031) | -0.707 (0.089) | $5.629 \times 10^{-13}$ | Pancreas | 1.079 (0.145) | $1.04 \times 10^{-13}$ |
| rs10908837 | *IL6R* | cg02856953 | -0.303 (0.039) | -0.120 (0.020) | $4.171 \times 10^{-9}$ | Whole Blood | 0.396 (0.083) | $2.05 \times 10^{-6}$ |

\* indicates surrogate variant used ( $r^2 > 0.8$ ), SNP – Single Nucleotide Polymorphism, Gene – likely implicated gene, CpG – 450K probe ID, CpG effect – effect estimate of SNP on methylation, eQTL effect – effect estimate of SNP on expression based on GTEx data, eQTL P – P-value for eQTL from GTEx, eQTL tissue – tissue type for observed effect according to GTEx, 2SMR effect – effect estimates from 2-Sample MR analysis, P-value – P-value for observed 2SMR effect

### Investigating the role of DNA methylation with gene expression and histone modification

To further dissect the relationship between DNA methylation and complex traits we sought to evaluate the influence of the methylation levels on local gene expression. We observed evidence of a causal relationship between methylation and expression at 8 of the 10 loci using data from the GTEx consortium (Table 4). Effect estimates suggest an inverse relationship (i.e. increased methylation results in decreased gene expression) at the *ADIPOQ* (low frequency signal) and *APOA1* loci, whereas a direct relationship was observed at the other 6 loci (i.e. increased methylation results in increased gene expression). We were unable to identify a surrogate variant (r^2^ > 0.8) to obtain a suitable effect estimate at the *LEPR* and *ADIPOQ* (common signal) loci.

mQTL at the *APOA1* and *IL6R* loci were also in high LD with previously reported histone quantitative trait loci (hQTL) based on findings by Grubert et al^44^. Results from our 2SMR analyses to evaluate the influence of methylation levels on histone modification provided strong evidence of a causal effect as well as an inverse relationship in each instance (Supplementary Table 6).

### Functional informatics

To better understand the functional role underlying these putative causal associations, variants which were ranked as the lead SNP based on the bivariate fine mapping analysis (using effect estimates on DNA methylation) were evaluated using VEP. Variants at the *ADIPOQ, ABO, LEPR, IL6R* and *APOA1* loci are located within histone mark peaks in adipose tissue according to data from the Roadmap Epigenomics Project and variants at *APOA1* & *CELSR2* reside within promoter flanking regions (Supplementary Table 7).

Every associated CpG site identified in this study resides within multiple histone mark peaks based on adipose tissue data from the Roadmap Epigenomics Project (Supplementary Table 8). All sites also reside in either enhancer or promoter regions with the exception of the CpG site near *ADIPOQ* (Supplementary Table 9). There was strong evidence of enrichment for regulatory annotations for both SNPs and CpG sites which supports previous evidence that they are likely to have a causal downstream effect on phenotypic variation (Supplementary Table 10).

## Discussion

We have designed a framework to evaluate the causal influences of DNA methylation on complex traits and disease using Mendelian randomization. For observed effects on cardiovascular traits that appear to be caused by methylation, we used bivariate fine mapping and joint likelihood mapping to evaluate if the putative causal variant influencing methylation was the same causal variant responsible for influencing the trait. This approach provides compelling evidence that cardiovascular traits are influenced by altered DNA methylation levels at the following 5 genes; *ABO*, *ADCY3*, *ADIPOQ*, *APOA1* and *IL6R*. Furthermore, two sample MR analyses provided evidence that DNA methylation levels influenced gene expression at these loci, suggesting that predicted functional effects for the causal variants indicate a coordinated system of effects that are consistent with causality. This was important to demonstrate, as single valid instruments meant that we were unable to robustly show that variants were not influencing methylation and traits independently. This type of approach is particularly attractive for therapeutic evaluation of drug targets as it can provide valuable insight into the underlying mechanisms between genetic variants and disease.

The *ABO* locus identified in this study has been associated with many different traits and diseases by previous studies^24; 51; 52^, as well as evidence implicating expression quantitative trait loci as putative causal SNPs for this effect ^53^. Here we provide evidence that DNA methylation may reside along the causal pathway to these observed effects (MR effect estimate: 0.29 (Standard error=0.06) change in trait per standard deviation change in methylation). A deletion (rs200533593) was found to be the putative causal variant for both the observed effect on DNA methylation and phenotypic variation.

The observed effect of genetic variation at *ADCY3* on body mass index is a relatively new finding^54-56^. In this study, our bivariate fine mapping analysis suggests that an intergenic variant (rs6737082) may be responsible for the observed signal which is mediated through DNA methylation at this locus (MR effect estimate: 0.11 (0.05)). Furthermore, a variant in LD with rs6737082 (rs713586, r^2^=0.80) has been previously reported to regulate DNA methylation at this location in adipose tissue^7^.

There were two independent effects detected in our study near the *ADIPOQ* gene which were associated with adiponectin. The common variant signal was located upstream of *ADIPOQ* within the *RFC4* gene but associated with DNA methylation levels proximal to *ADIPOQ*, which can help explain this variant’s observed effect on adiponectin (MR effect estimate: -0.36 (0.12)). Concordance in the bivariate fine mapping analysis suggested a non-coding transcript variant (rs169109) was responsible. The lead SNP from the ADIPOGen consortium^57^ at this locus (rs6810075) is neither an mQTL nor in high LD with rs169109 (r^2^ = 0.20), suggesting that these two association signals are influencing adiponectin levels by alternative biological mechanisms. The low frequency variant signal was previously detected by the UK10K project ^33^, although bivariate fine mapping results as this locus suggest that the causal mQTL was in linkage disequilibrium with the trait-associated variant.

The CpG site associated with the mQTL at this locus resides between the *APOA1* gene and *APOA1-antisense* (*APOA1-AS*), a negative transcriptional regulatory of *APOA1* which has been shown to increase *APOA1* expression both in vitro and in vivo^58^. The highest ranked mQTL based on our bivariate fine mapping using estimates with DNA methylation is in a promoter region upstream of *APOA1*, suggesting that it may be more likely influencing *APOA1* rather than *APOA1-AS*. There are previously reported GWAS association signals at this locus with lipid traits^59; 60^. However, given the evidence in this study of a causal effect with DNA methylation (MR effect estimate: -0.30 (0.08)) it is likely that these are downstream effects of the observed effect on Apo A1 variation. Furthermore, it is more biologically plausible this is the case as this gene is responsible for the protein synthesis of Apo A1.

The signal at the *IL6R* locus has been previously associated with a range of traits related to respiratory and cardiovascular health ^61-63^. Our results suggest that genetic variation at *IL6R* influences DNA methylation at this region, which in turn will have a downstream effect on interleukin-6 and subsequently other traits and diseases (MR effect estimate: 0.47 (0.18)). Furthermore, this association signal was not in LD with a previously reported missense variant at this locus (rs2228145, r^2^ = 0.47 in ALSPAC) which was also supported by findings from an in-depth functional study of this variant ^64^.

Evidence from the GTEx consortium suggests that mQTL at all eight of the loci with available expression data overlap with eQTL effects, which serves as an independent replication of the relationships discovered through DNA methylation levels. It is biologically plausible that a variant’s impact of DNA methylation levels may have a downstream effect on gene expression along the causal pathway to disease^65; 66^, which may help explain these observations. Effects at four loci in particular appear to be biologically plausible in this regard, as the likely genes influenced by these variants are involved in the protein synthesis of the associated trait (i.e. *ADIPOQ* with adiponectin, *APOB* with Apo B, *APOAI* with Apo A1 and *IL6R* with interleukin-6). Furthermore, each CpG site identified in this study resides within histone mark peaks in adiposity tissue according to data from the Roadmap Epigenomics project where we observed enrichment compared to matched CpG sites in similar regions of the genome.

As with any study which applies MR using a single instrument to investigate causal relationships in epidemiology, an important limitation is the inability to disentangle potential pleiotropic effects where the same causal variant influences both exposure (i.e. DNA methylation) and outcome (i.e. cardiovascular trait) through independent pathways. To reduce the possibility of this we selected mQTL in our study that were only influencing proximal CpG sites and not elsewhere in the epigenome, as such instruments would be more likely to influencing traits via alternative biological mechanisms. Future studies which continue to uncover mQTL across the genome (as well as across various tissue types) should facilitate analyses which are able to robustly address concerns over pleiotropy by using methods such as MR-Egger ^67^.

Weak instrumental variables and reverse causation are other factors which can bias MR analyses. Our analysis is unlikely to have suffered from the former as each mQTL had a large effect on DNA methylation in cis (P < 1.0×10^-14^) and were robustly associated with traits which we were able to replicate using results from studies with large population samples. We conducted analyses to evaluate whether reverse causation was an issue in our study (i.e. trait variation was causal to changes in DNA methylation at each locus), although results suggested that this was not the case.

In this study we have demonstrated the value of two-sample Mendelian randomization (2SMR) to undertake MR analyses using summary statistics ^39; 68^. This allowed us to provide evidence of replication for the observed effects in our study as well as investigate the relationship between DNA methylation and expression along the causal pathway to disease. This approach has the attractive advantage of enabling the potential epigenetic-complex trait interplay to be interrogated on a much wider scale, foregoing the requirement that ‘omic’ data and phenotypes are measured in the same sample.

## Description of Supplemental Data

Supplemental Data include one descriptive methods section for phenotyping and 10 tables.

## Acknowledgements

We are extremely grateful to all the families who took part in this study, the midwives for their help in recruiting them, and the whole ALSPAC team, which includes interviewers, computer and laboratory technicians, clerical workers, research scientists, volunteers, managers, receptionists and nurses. The UK Medical Research Council and the Wellcome Trust (Grant ref: 102215/2/13/2) and the University of Bristol provide core support for ALSPAC. GWAS data was generated by Sample Logistics and Genotyping Facilities at the Wellcome Trust Sanger Institute and LabCorp (Laboratory Corporation of America) using support from 23andMe. Methylation data in the ALSPAC cohort was generated as part of the UK BBSRC funded (BB/I025751/1 and BB/I025263/1) Accessible Resource for Integrated Epigenomic Studies (ARIES, http://www.ariesepigenomics.org.uk).

This publication is the work of the authors and Tom G Richardson will serve as guarantor for the contents of this paper. This work was supported by the UK Medical Research Council (MRC Integrative Epidemiology Unit, MC UU 12013/1, MC UU 12013/2, MC UU 12013/3, MC UU 12013/8).

## Web Resources

ARIES explorer - www.ariesepigenomics.org.uk/

GWAS catalog - https://www.ebi.ac.uk/gwas/

GTEx - www.gtexportal.org/

Ensembl - www.ensembl.org/

Epigenomics Roadmap Project - www.roadmapepigenomics.org/

snpSNAP - https://data.broadinstitute.org/mpg/snpsnap/

## References

1. Edwards, S.L., Beesley, J., French, J.D., and Dunning, A.M. (2013). Beyond GWASs: illuminating the dark road from association to function. American journal of human genetics 93, 779–797.

2. Zhu, Z., Zhang, F., Hu, H., Bakshi, A., Robinson, M.R., Powell, J.E., Montgomery, G.W., Goddard, M.E., Wray, N.R., Visscher, P.M., et al. (2016). Integration of summary data from GWAS and eQTL studies predicts complex trait gene targets. Nat Genet 48, 481–487.

3. Burkhardt, R., Kirsten, H., Beutner, F., Holdt, L.M., Gross, A., Teren, A., Tonjes, A., Becker, S., Krohn, K., Kovacs, P., et al. (2015). Integration of Genome-Wide SNP Data and Gene-Expression Profiles Reveals Six Novel Loci and Regulatory Mechanisms for Amino Acids and Acylcarnitines in Whole Blood. PLoS genetics 11, e1005510.

4. Pavlides, J.M., Zhu, Z., Gratten, J., McRae, A.F., Wray, N.R., and Yang, J. (2016). Predicting gene targets from integrative analyses of summary data from GWAS and eQTL studies for 28 human complex traits. Genome medicine 8, 84.

5. Mancuso, N., Shi, H., Goddard, P., Kichaev, G., Gusev, A., and Pasaniuc, B. (2017). Integrating Gene Expression with Summary Association Statistics to Identify Genes Associated with 30 Complex Traits. American journal of human genetics 100, 473–487.

6. Kulis, M., Heath, S., Bibikova, M., Queiros, A.C., Navarro, A., Clot, G., Martinez-Trillos, A., Castellano, G., Brun-Heath, I., Pinyol, M., et al. (2012). Epigenomic analysis detects widespread gene-body DNA hypomethylation in chronic lymphocytic leukemia. Nat Genet 44, 1236–1242.

7. Grundberg, E., Meduri, E., Sandling, J.K., Hedman, A.K., Keildson, S., Buil, A., Busche, S., Yuan, W., Nisbet, J., Sekowska, M., et al. (2013). Global analysis of DNA methylation variation in adipose tissue from twins reveals links to disease-associated variants in distal regulatory elements. American journal of human genetics 93, 876–890.

8. Gaunt, T.R., Shihab, H.A., Hemani, G., Min, J.L., Woodward, G., Lyttleton, O., Zheng, J., Duggirala, A., McArdle, W.L., Ho, K., et al. (2016). Systematic identification of genetic influences on methylation across the human life course. Genome biology 17, 61.

9. Shi, J., Marconett, C.N., Duan, J., Hyland, P.L., Li, P., Wang, Z., Wheeler, W., Zhou, B., Campan, M., Lee, D.S., et al. (2014). Characterizing the genetic basis of methylome diversity in histologically normal human lung tissue. Nature communications 5, 3365.

10. Bell, J.T., Tsai, P.C., Yang, T.P., Pidsley, R., Nisbet, J., Glass, D., Mangino, M., Zhai, G., Zhang, F., Valdes, A., et al. (2012). Epigenome-wide scans identify differentially methylated regions for age and age-related phenotypes in a healthy ageing population. PLoS genetics 8, e1002629.

11. Wahl, S., Drong, A., Lehne, B., Loh, M., Scott, W.R., Kunze, S., Tsai, P.C., Ried, J.S., Zhang, W., Yang, Y., et al. (2017). Epigenome-wide association study of body mass index, and the adverse outcomes of adiposity. Nature 541, 81–86.

12. Liang, L., Willis-Owen, S.A., Laprise, C., Wong, K.C., Davies, G.A., Hudson, T.J., Binia, A., Hopkin, J.M., Yang, I.V., Grundberg, E., et al. (2015). An epigenome-wide association study of total serum immunoglobulin E concentration. Nature 520, 670–674.

13. Gusev, A., Ko, A., Shi, H., Bhatia, G., Chung, W., Penninx, B.W., Jansen, R., de Geus, E.J., Boomsma, D.I., Wright, F.A., et al. (2016). Integrative approaches for large-scale transcriptome-wide association studies. Nat Genet 48, 245–252.

14. Powell, J.E., Fung, J.N., Shakhbazov, K., Sapkota, Y., Cloonan, N., Hemani, G., Hillman, K.M., Kaufmann, S., Luong, H.T., Bowdler, L., et al. (2016). Endometriosis risk alleles at 1p36.12 act through inverse regulation of CDC42 and LINC00339. Human molecular genetics 25, 5046–5058.

15. Rawlik, K., Rowlatt, A., and Tenesa, A. (2016). Imputation of DNA Methylation Levels in the Brain Implicates a Risk Factor for Parkinson’s Disease. Genetics 204, 771–781.

16. Davey Smith, G., and Hemani, G. (2014). Mendelian randomization: genetic anchors for causal inference in epidemiological studies. Human molecular genetics 23, R89–98.

17. Davey Smith, G., and Ebrahim, S. (2003). ‘Mendelian randomization’: can genetic epidemiology contribute to understanding environmental determinants of disease? International journal of epidemiology 32, 1–22.

18. Relton, C.L., and Davey Smith, G. (2012). Two-step epigenetic Mendelian randomization: a strategy for establishing the causal role of epigenetic processes in pathways to disease. International journal of epidemiology 41, 161–176.

19. Millstein, J., Zhang, B., Zhu, J., and Schadt, E.E. (2009). Disentangling molecular relationships with a causal inference test. BMC genetics 10, 23.

20. Richmond, R.C., Hemani, G., Tilling, K., Davey Smith, G., and Relton, C.L. (2016). Challenges and novel approaches for investigating molecular mediation. Human molecular genetics 25, R149–R156.

21. Ference, B.A., Yoo, W., Alesh, I., Mahajan, N., Mirowska, K.K., Mewada, A., Kahn, J., Afonso, L., Williams, K.A., Sr., and Flack, J.M. (2012). Effect of long-term exposure to lower low-density lipoprotein cholesterol beginning early in life on the risk of coronary heart disease: a Mendelian randomization analysis. Journal of the American College of Cardiology 60, 2631–2639.

22. Vimaleswaran, K.S., Berry, D.J., Lu, C., Tikkanen, E., Pilz, S., Hiraki, L.T., Cooper, J.D., Dastani, Z., Li, R., Houston, D.K., et al. (2013). Causal relationship between obesity and vitamin D status: bi-directional Mendelian randomization analysis of multiple cohorts. PLoS medicine 10, e1001383.

23. Chun, S., Casparino, A., Patsopoulos, N.A., Croteau-Chonka, D.C., Raby, B.A., De Jager, P.L., Sunyaev, S.R., and Cotsapas, C. (2017). Limited statistical evidence for shared genetic effects of eQTLs and autoimmune-disease-associated loci in three major immune-cell types. Nat Genet 49, 600–605.

24. Pickrell, J.K., Berisa, T., Liu, J.Z., Segurel, L., Tung, J.Y., and Hinds, D.A. (2016). Detection and interpretation of shared genetic influences on 42 human traits. Nat Genet 48, 709–717.

25. Giambartolomei, C., Vukcevic, D., Schadt, E.E., Franke, L., Hingorani, A.D., Wallace, C., and Plagnol, V. (2014). Bayesian test for colocalisation between pairs of genetic association studies using summary statistics. PLoS genetics 10, e1004383.

26. Hemani, G., Tilling, K., and Davey Smith, G. (2017). Orienting The Causal Relationship Between Imprecisely Measured Traits Using Genetic Instruments.

27. Boyd, A., Golding, J., Macleod, J., Lawlor, D.A., Fraser, A., Henderson, J., Molloy, L., Ness, A., Ring, S., and Davey Smith, G. (2013). Cohort Profile: the ‘children of the 90s’--the index offspring of the Avon Longitudinal Study of Parents and Children. International journal of epidemiology 42, 111–127.

28. Fraser, A., Macdonald-Wallis, C., Tilling, K., Boyd, A., Golding, J., Davey Smith, G., Henderson, J., Macleod, J., Molloy, L., Ness, A., et al. (2013). Cohort Profile: the Avon Longitudinal Study of Parents and Children: ALSPAC mothers cohort. International journal of epidemiology 42, 97–110.

29. Relton, C.L., Gaunt, T., McArdle, W., Ho, K., Duggirala, A., Shihab, H., Woodward, G., Lyttleton, O., Evans, D.M., Reik, W., et al. (2015). Data Resource Profile: Accessible Resource for Integrated Epigenomic Studies (ARIES). International journal of epidemiology.

30. Touleimat, N., and Tost, J. (2012). Complete pipeline for Infinium((r)) Human Methylation 450K BeadChip data processing using subset quantile normalization for accurate DNA methylation estimation. Epigenomics 4, 325–341.

31. Pidsley, R., Y Wong, C.C., Volta, M., Lunnon, K., Mill, J., and Schalkwyk, L.C. (2013). A data-driven approach to preprocessing Illumina 450K methylation array data. BMC Genomics 14, 293.

32. Jaffe, A.E., and Irizarry, R.A. (2014). Accounting for cellular heterogeneity is critical in epigenome-wide association studies. Genome biology 15, R31.

33. The UK10K Consortium. (2015). The UK10K project identifies rare variants in health and disease. Nature.

34. Howie, B.N., Donnelly, P., and Marchini, J. (2009). A flexible and accurate genotype imputation method for the next generation of genome-wide association studies. PLoS genetics 5, e1000529.

35. Naeem, H., Wong, N.C., Chatterton, Z., Hong, M.K., Pedersen, J.S., Corcoran, N.M., Hovens, C.M., and Macintyre, G. (2014). Reducing the risk of false discovery enabling identification of biologically significant genome-wide methylation status using the HumanMethylation450 array. BMC genomics 15, 51.

36. Yang, J., Ferreira, T., Morris, A.P., Medland, S.E., Genetic Investigation of, A.T.C., Replication, D.I.G., Meta-analysis, C., Madden, P.A., Heath, A.C., Martin, N.G., et al. (2012). Conditional and joint multiple-SNP analysis of GWAS summary statistics identifies additional variants influencing complex traits. Nat Genet 44, 369-375, S361–363.

37. Turner, S.D. (2014). qqman: an R package for visualizing GWAS results using Q-Q and manhattan plots.

38. Henningsen, A., Hamann, J.D. (2007). systemfit: A Package for Estimating Systems of Simultaneous Equations in R. Journal of Statistical Software 23, 1–40.

39. Burgess, S., Scott, R.A., Timpson, N.J., Davey Smith, G., Thompson, S.G., and Consortium, E.-I. (2015). Using published data in Mendelian randomization: a blueprint for efficient identification of causal risk factors. Eur J Epidemiol 30, 543–552.

40. MacArthur, J., Bowler, E., Cerezo, M., Gil, L., Hall, P., Hastings, E., Junkins, H., McMahon, A., Milano, A., Morales, J., et al. (2017). The new NHGRI-EBI Catalog of published genome-wide association studies (GWAS Catalog). Nucleic acids research 45, D896–D901.

41. Naitza, S., Porcu, E., Steri, M., Taub, D.D., Mulas, A., Xiao, X., Strait, J., Dei, M., Lai, S., Busonero, F., et al. (2012). A genome-wide association scan on the levels of markers of inflammation in Sardinians reveals associations that underpin its complex regulation. PLoS genetics 8, e1002480.

42. Benner, C., Spencer, C.C., Havulinna, A.S., Salomaa, V., Ripatti, S., and Pirinen, M. (2016). FINEMAP: efficient variable selection using summary data from genome-wide association studies. Bioinformatics 32, 1493–1501.

43. Consortium, G.T. (2013). The Genotype-Tissue Expression (GTEx) project. Nat Genet 45, 580–585.

44. Grubert, F., Zaugg, J.B., Kasowski, M., Ursu, O., Spacek, D.V., Martin, A.R., Greenside, P., Srivas, R., Phanstiel, D.H., Pekowska, A., et al. (2015). Genetic Control of Chromatin States in Humans Involves Local and Distal Chromosomal Interactions. Cell 162, 1051–1065.

45. Rose, N.R., and Klose, R.J. (2014). Understanding the relationship between DNA methylation and histone lysine methylation. Biochimica et biophysica acta 1839, 1362–1372.

46. Liu, L., Jin, G., and Zhou, X. (2015). Modeling the relationship of epigenetic modifications to transcription factor binding. Nucleic acids research 43, 3873–3885.

47. McLaren, W., Gil, L., Hunt, S.E., Riat, H.S., Ritchie, G.R., Thormann, A., Flicek, P., and Cunningham, F. (2016). The Ensembl Variant Effect Predictor. Genome biology 17, 122.

48. Yates, A., Akanni, W., Amode, M.R., Barrell, D., Billis, K., Carvalho-Silva, D., Cummins, C., Clapham, P., Fitzgerald, S., Gil, L., et al. (2016). Ensembl 2016. Nucleic acids research 44, D710–716.

49. Bernstein, B.E., Stamatoyannopoulos, J.A., Costello, J.F., Ren, B., Milosavljevic, A., Meissner, A., Kellis, M., Marra, M.A., Beaudet, A.L., Ecker, J.R., et al. (2010). The NIH Roadmap Epigenomics Mapping Consortium. Nature biotechnology 28, 1045–1048.

50. Pers, T.H., Timshel, P., and Hirschhorn, J.N. (2015). SNPsnap: a Web-based tool for identification and annotation of matched SNPs. Bioinformatics 31, 418–420.

51. Nikpay, M., Goel, A., Won, H.H., Hall, L.M., Willenborg, C., Kanoni, S., Saleheen, D., Kyriakou, T., Nelson, C.P., Hopewell, J.C., et al. (2015). A comprehensive 1,000 Genomes-based genome-wide association meta-analysis of coronary artery disease. Nat Genet 47, 1121–1130.

52. Hinds, D.A., Buil, A., Ziemek, D., Martinez-Perez, A., Malik, R., Folkersen, L., Germain, M., Malarstig, A., Brown, A., Soria, J.M., et al. (2016). Genome-wide association analysis of self-reported events in 6135 individuals and 252 827 controls identifies 8 loci associated with thrombosis. Human molecular genetics 25, 1867–1874.

53. Wessel, J., Chu, A.Y., Willems, S.M., Wang, S., Yaghootkar, H., Brody, J.A., Dauriz, M., Hivert, M.F., Raghavan, S., Lipovich, L., et al. (2015). Low-frequency and rare exome chip variants associate with fasting glucose and type 2 diabetes susceptibility. Nature communications 6, 5897.

54. Locke, A.E., Kahali, B., Berndt, S.I., Justice, A.E., Pers, T.H., Day, F.R., Powell, C., Vedantam, S., Buchkovich, M.L., Yang, J., et al. (2015). Genetic studies of body mass index yield new insights for obesity biology. Nature 518, 197–206.

55. Warrington, N.M., Howe, L.D., Paternoster, L., Kaakinen, M., Herrala, S., Huikari, V., Wu, Y.Y., Kemp, J.P., Timpson, N.J., St Pourcain, B., et al. (2015). A genome-wide association study of body mass index across early life and childhood. International journal of epidemiology 44, 700–712.

56. Felix, J.F., Bradfield, J.P., Monnereau, C., van der Valk, R.J., Stergiakouli, E., Chesi, A., Gaillard, R., Feenstra, B., Thiering, E., Kreiner-Moller, E., et al. (2016). Genome-wide association analysis identifies three new susceptibility loci for childhood body mass index. Human molecular genetics 25, 389–403.

57. Dastani, Z., Hivert, M.F., Timpson, N., Perry, J.R., Yuan, X., Scott, R.A., Henneman, P., Heid, I.M., Kizer, J.R., Lyytikainen, L.P., et al. (2012). Novel loci for adiponectin levels and their influence on type 2 diabetes and metabolic traits: a multi-ethnic meta-analysis of 45,891 individuals. PLoS genetics 8, e1002607.

58. Halley, P., Kadakkuzha, B.M., Faghihi, M.A., Magistri, M., Zeier, Z., Khorkova, O., Coito, C., Hsiao, J., Lawrence, M., and Wahlestedt, C. (2014). Regulation of the apolipoprotein gene cluster by a long noncoding RNA. Cell Rep 6, 222–230.

59. Lu, X., Huang, J., Mo, Z., He, J., Wang, L., Yang, X., Tan, A., Chen, S., Chen, J., Gu, C.C., et al. (2016). Genetic Susceptibility to Lipid Levels and Lipid Change Over Time and Risk of Incident Hyperlipidemia in Chinese Populations. Circulation Cardiovascular genetics 9, 37–44.

60. Kurano, M., Tsukamoto, K., Kamitsuji, S., Kamatani, N., Hara, M., Ishikawa, T., Kim, B.J., Moon, S., Jin Kim, Y., and Teramoto, T. (2016). Genome-wide association study of serum lipids confirms previously reported associations as well as new associations of common SNPs within PCSK7 gene with triglyceride. Journal of human genetics 61, 427–433.

61. Ferreira, M.A., Matheson, M.C., Duffy, D.L., Marks, G.B., Hui, J., Le Souef, P., Danoy, P., Baltic, S., Nyholt, D.R., Jenkins, M., et al. (2011). Identification of IL6R and chromosome 11q13.5 as risk loci for asthma. Lancet 378, 1006–1014.

62. Dehghan, A., Dupuis, J., Barbalic, M., Bis, J.C., Eiriksdottir, G., Lu, C., Pellikka, N., Wallaschofski, H., Kettunen, J., Henneman, P., et al. (2011). Meta-analysis of genome-wide association studies in >80 000 subjects identifies multiple loci for C-reactive protein levels. Circulation 123, 731–738.

63. Khera, A.V., Emdin, C.A., Drake, I., Natarajan, P., Bick, A.G., Cook, N.R., Chasman, D.I., Baber, U., Mehran, R., Rader, D.J., et al. (2016). Genetic Risk, Adherence to a Healthy Lifestyle, and Coronary Disease. The New England journal of medicine 375, 2349–2358.

64. van Dongen, J., Jansen, R., Smit, D., Hottenga, J.J., Mbarek, H., Willemsen, G., Kluft, C., Collaborators, A., Penninx, B.W., Ferreira, M.A., et al. (2014). The contribution of the functional IL6R polymorphism rs2228145, eQTLs and other genome-wide SNPs to the heritability of plasma sIL-6R levels. Behavior genetics 44, 368–382.

65. Jones, P.A., and Takai, D. (2001). The role of DNA methylation in mammalian epigenetics. Science 293, 1068–1070.

66. Baylin, S.B., Esteller, M., Rountree, M.R., Bachman, K.E., Schuebel, K., and Herman, J.G. (2001). Aberrant patterns of DNA methylation, chromatin formation and gene expression in cancer. Human molecular genetics 10, 687–692.

67. Bowden, J., Davey Smith, G., and Burgess, S. (2015). Mendelian randomization with invalid instruments: effect estimation and bias detection through Egger regression. International journal of epidemiology 44, 512–525.

68. Hemani, G., Zheng, J., Wade, K.H., Laurin, C., Elsworth, E., Burgess, S., Bowden, J., Langdon, R., Tan, V., Yarmolinsky, J., et al. (2016). MR-Base: a platform for systematic causal inference across the phenome using billions of genetic associations.

